# Rationally designed split Lettuce aptamer based on large scale mutational analysis

**DOI:** 10.64898/2026.07.10.737091

**Authors:** Alexandra M. Adams, Edward B. Pimentel, N. Duane Loh, Yasser Gidi, Linus A. Hein, Michael Eisenstein, H. Tom Soh

## Abstract

Split aptamer biosensors offer exceptionally low background by assembling only in the presence of a target analyte; however, their performance is frequently limited by the lack of robust design rules for selecting effective split sites. Existing approaches largely rely on heuristic, structure-based assumptions that are poorly validated and often yield suboptimal signal. Herein, we introduce a systematic, data-driven strategy for identifying high-performance split sites within fluorogenic DNA aptamers. Using our massively-parallel aptamer performance analyzer (MAPA) platform, we performed comprehensive single- and double-mutant analysis of the DFAME-binding region of the fluorogenic DNA aptamer Lettuce, informed by its three-dimensional structure. Dimensionality reduction and clustering of the resulting sequence-function landscape revealed mutation-tolerant elements within the binding domain that are suitable for splitting while preserving fluorophore activation. Sensors constructed using these non-intuitive split sites, which are unconventional by standard design principles, exhibited a nearly four-fold improvement in fluorescence signal-to-background ratio for SARS-CoV-2 RNA detection compared to a prior split-Lettuce design. The same split architecture also enabled robust detection of high-pathogenicity H5Nx avian influenza RNA. These results demonstrate that large-scale, data-driven interrogation of aptamer sequence-function relationships can identify non-intuitive split sites and provide a proof-of-concept framework for developing measurement-based design principles for split-aptamer biosensors.

## Background

Split aptamer sensors are a class of nucleic-acid-based biorecognition elements that have proven valuable for small molecule and protein detection in a variety of applications. These sensors are generated by dividing a DNA or RNA aptamer into two or more short segments^1^. Individually, these split-aptamer segments remain largely unstructured and inactive, but they retain the capacity to reassemble into a fully-folded and functional aptamer structure in the presence of their target ligand^2^. This selective refolding based on specific target recognition events makes split aptamers useful for biosensing and diagnostics due to the low background signal produced in the absence of target. Such sensors can be engineered to generate a signal based on assembly-induced changes in the relative positioning of functional tags attached to each individual fragment^3–5^, such as Förster resonance energy transfer (FRET)-compatible fluorophore pairs^6,7^. Alternatively, several groups have developed fluorogenic ‘green fluorescent protein (GFP) mimic’ aptamers such as Lettuce or Broccoli^8–10^, which only generate a fluorescence signal when their fully folded confirmations bind to their paired fluorophore. Lettuce binds to chromophores including DFHBI-1T, DFHO, and DFAME, which normally have low background fluorescence but exhibit dramatically enhanced fluorescence when bound to the GFP mimic aptamer. When the split Lettuce aptamer sensor recognizes its complementary nucleic acid target sequence, the split aptamer reassembles and is subsequently able to bind to one of these chromophores to produce a fluorescent readout. In prior work, Lettuce has been successfully developed into split aptamer sensors for SARS-CoV-2 RNA^9^ as well as Hepatitis B viral DNA^8^.

It remains challenging to identify optimal split sites within a given aptamer that enable high-performance biosensing^11,12^. Ideally, the split aptamer fragments should be incapable of binding the target individually or to each other in the absence of target, with efficient reassembly into the fully folded aptamer structure only occurring in the presence of both fragments and the target.

Unfortunately, target-independent assembly is still possible with a suboptimal split site, resulting in high background signal and reducing the sensitivity of the biosensor. Alternately, suboptimal split sites also result in fragments that cannot fully reconstitute the aptamer structure, yielding an ineffective sensor. Indeed, the Lettuce sensors described above only achieved modest increases in fluorescence upon binding their target, limiting their utility as viral sensors. Previous split aptamer generation efforts have relied on rational design principles coupled with trial-and-error optimization processes, but these approaches do not necessarily offer a straightforward solution. The 2D aptamer structures generated by conventional prediction algorithms are generally best suited for identifying stem-loops within the sequence. Stem-forming regions are generally thought to not be involved in target binding, but instead to contribute primarily to the complex 3D folding of the full construct. As such, most rational design efforts have focused on splitting aptamers at these stem-forming regions, with the goal of disrupting folding in the absence of target while still preserving the ability to bind target. Most design efforts also avoid splitting at sites close to target-binding nucleotides, which are typically assumed to reside within large loops. However, Zhou and colleagues have demonstrated that some loop regions considered to be functionally essential could nevertheless tolerate splitting and enable downstream biosensing^13^. This demonstrates that the field still lacks a clear understanding of which aptamer features offer optimal sites for split-aptamer sensor generation.

In this work, we show that high-throughput mutational analysis can be coupled with detailed structural models to identify counter-intuitive but effective split sites for the Lettuce aptamer, which we subsequently employed to design a greatly improved SARS-CoV-2 split-aptamer sensor and novel avian influenza split-aptamer sensor. We relied on insights from a recently-derived set of 3D structures of Lettuce in complex with the chromophores DFHBI-1T, DFHO, and DFAME, which provided novel mechanistic insights into the performance of the aptamer^14^ and revealed inaccuracies in previously-predicted 2D structural models. High-throughput mutational analysis provides important sequence-function data that can be used to guide aptamer selection and engineering^15–18^, and we recently developed a platform for performing such experiments called the massively-parallel aptamer performance analyzer (MAPA)^19^. The MAPA platform builds upon our previous work of using a benchtop Illumina MiSeq flow cell for mapping sequence and function relationships directly^18^. By combining mutational analysis with functional analysis, we identified nucleotides which were not critical for chromophore binding to Lettuce. We hypothesized that nucleotides that tolerate mutation without disrupting Lettuce function would be more likely to tolerate the larger perturbation introduced by a split site, provided that target-induced proximity allows the aptamer to refold productively. We therefore generated an extensive library of single and double mutants of Lettuce based on insights from these crystal structures, which we then screened using MAPA. We employed a combination of vector-quantized clustering and dimensionality-reducing algorithms to identify sequence/structure elements that can be safely modified without impairing aptamer-chromophore binding, indicating features that are likely to be amenable to the introduction of split sites as well. We then characterized several of the most promising split sites experimentally by developing a SARS-CoV-2 split-aptamer sensor; all three performed well, and one construct achieved a nearly four-fold improvement in fluorescence signal-to-background ratio compared to the previously published split-Lettuce sensor for this target. We subsequently created a novel sensor based on the same split aptamer architecture that could detect RNA from high-pathogenicity (HP) H5Nx avian influenza variants^20^, supporting that our split architecture can be successfully applied to another RNA target. These results demonstrate that systematic data-driven analysis can identify optimal split sites within aptamer sequences that would be far more challenging to identify—and perhaps even overlooked entirely—with more laborious approaches based on human intuition and trial and error. Together, these results demonstrate that high-throughput sequence–function analysis can identify split sites within fluorogenic DNA aptamers that would be difficult to predict by human intuition or trial-and-error alone. More broadly, this work provides a proof-of-concept framework for developing measurement-based design principles for split-aptamer sensors

## EXPERIMENTAL SECTION

### Materials

The single and double mutant libraries were purchased from Twist Biosciences with sequencing adapters attached at the 3′ and 5′ ends of each sequence (Data link 1). All other DNA sequences were purchased from Integrated DNA Technologies (IDT) with standard desalting for unmodified oligos and HPLC purification for oligos with fluorophores and resuspended in ultra-pure water upon arrival (Table S1). RNA sequences were also ordered from IDT with standard desalting and resuspended in 1x IDTE buffer (Table S2). 1x IDTE buffer (pH 8.0) was also ordered from IDT. All other reagents and buffers were purchased from Thermo Fisher Scientific unless otherwise noted. DFAME was purchased from MedChem Express and resuspended in new 100% DMSO to a concentration of 25 mM. DFHBI-1T was purchased from Lucerna and resuspended in 100% DMSO to a concentration of 25 mM. EcoRI restriction enzyme (NEB #R3101T) was purchased from New England Biolabs. All solution-based experiments were performed in CoStar 96-well, half-area opaque plates and measured using a BioTek Synergy H1 microplate reader. All surface-coupled experiments were done with MyOne Dyna C1 streptavidin-coated 1-µm magnetic beads and measured using an Agilent Novocyte Quanteon flow cytometer. Sequencing was done using an Illumina MiSeq with all provided reagents in the Nextera Index Kit (#15055290) and 150-cycle MiSeq Reagent Kit v3 (#MS-102-3001) from Illumina. All full-construct experiments were done in 1x Lettuce buffer (40 mM HEPES, 120 mM KCl, 1 mM MgCl2), unless otherwise noted. All split-construct experiments were done in 1x high-Mg2+ Lettuce buffer (40 mM HEPES, 120 mM KCl, 5 mM MgCl2) unless otherwise noted.

### Mutant library generation

The mutation sites for the mutant library were chosen based on prior knowledge of bases outside of the stem regions. These bases are known to be involved in the tertiary folding and/or the binding of the target, whereas disruptions to stem features are known to completely unfold the aptamer. We ordered all single and double mutants for bases 7–28 and 43–47 (Fig. 1D) from Twist Biosciences as an oligo pool with sequencing adapters attached at the 5′ and 3′ ends. We also incorporated half of the EcoRI restriction enzyme site on the 3′ end of the library between the mutant library sequence and the sequencing adapter, so that we could remove the adapter after sequencing (Lettuce_Adap_EcoRI). We also incorporated a poly(T)10 linker between the 5′ adapter and our mutant library sequence to provide flexibility. Upon receiving the libraries, we resuspended the sequences in ultra-pure water to a concentration of 1.25 mg/mL based on Twist Bioscience’s protocol. We used PCR with Go-Taq Hot Start Colorless Master Mix polymerase to attach unique sequencing indexes to each library. The conditions of the PCR were: 95 °C for 120 s, up to eight cycles of 95 °C for 15 s, 54 °C for 15 s, and 72 °C for 30 s, followed by 72 °C for 60 s. We performed polyacrylamide gel electrophoresis (PAGE) with 10% Invitrogen Novex TBE Gels to purify the indexed libraries and confirm proper addition of the indexes to the libraries. We extracted the mutant library with the sequencing indices from the PAGE gel using QIAquick purification kits from Qiagen. We measured the library concentration on a Qubit fluorometer (Thermo Fisher Scientific) and diluted the purified library to 20 pM in water. We followed the guidelines from Illumina to mix our libraries with PhiX. We additionally added an in-house positive control pool known as fiducial mark sequences, as previously described^18^.

**Figure 1.**
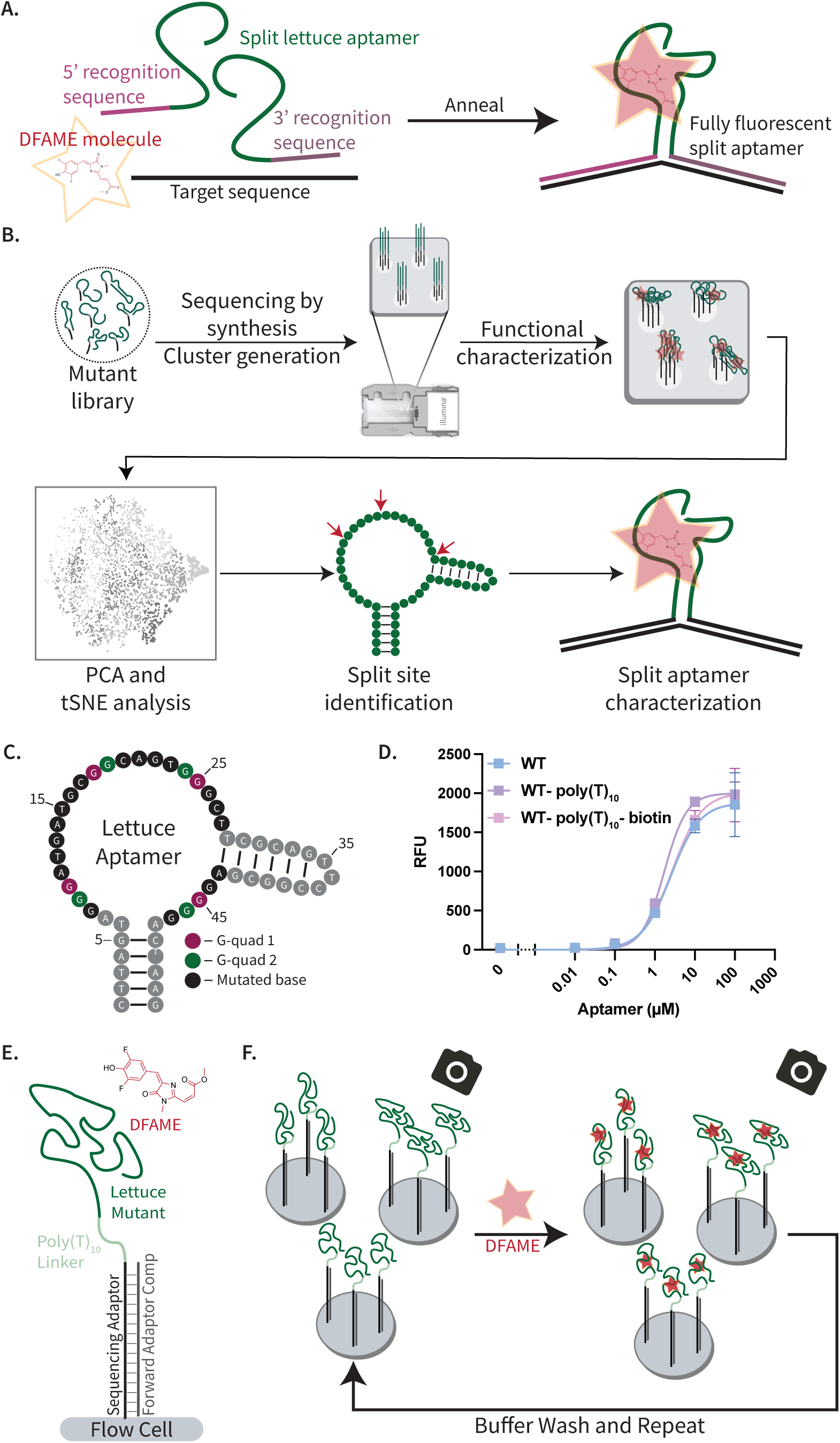
Lettuce aptamer mutational analysis workflow. A) Split aptamer schematic where two halves of the split aptamer are incubated with the target sequence and DFAME chromophore and are annealed to fold into the full construct and bind DFAME. B) We begin by generating and sequencing clusters from our mutant aptamer library on an Illumina flow cell, followed by functional characterization on the MAPA platform, computational analysis to identify promising split sites, and characterization of the resulting split aptamers. C) 2D structure of the Lettuce aptamer, with black and colored bases depicting mutated bases; the latter are participants in G-quadruplex structures. D) Binding curves (fitted with the Hill slope equation) for the wildtype (WT) sequence alone, with a poly(T)10 linker, and with a biotinylated poly(T)10 linker. N = 3, error bars represent the standard deviation. E) Labeled diagram of a sequence cluster on the flow cell surface with a complement binding the bases not involved in the Lettuce aptamer. F) The MAPA analysis process. First, a background image is taken with no DFAME present. The aptamer mutant clusters are then incubated with DFAME for 10 min and imaged again to identify mutants with enhanced fluorescence. The DFAME is removed by washing with buffer, and another background image is taken before the next concentration of DFAME is flowed in.

### Sequencing and MAPA characterization

Following standard Illumina MiSeq procedures, we loaded the cartridge with the pooled DNA libraries and controls. We then performed sequencing as per standard Illumina protocols. We downloaded the FastQ files from the MiSeq for subsequent data processing. We then utilized that same flow cell on our MAPA platform^19^. We followed all preparation procedures described in Gidi *et al.* (2025) for MAPA, unless otherwise noted. Briefly, we first incubated with a complement containing the other half of the restriction site (EcoRI comp) and the EcoRI restriction enzyme to remove the 3′ reverse adapter and sequencing artifacts at 37 °C for 15 min. We performed three rounds of digestion to confirm that all reverse adapters had been fully removed. We then prepared the flow cell for MAPA via a 100% formamide wash and 1x Lettuce buffer incubation. Next, we annealed and incubated with a forward adapter complement labeled with Cy3 (FAC-Cy3) so that we could image each cluster and identify its relative size. We then removed the FAC-Cy3 strand using another formamide wash and added unlabeled FAC, complementary sequences for the lawn primers (LC1, LC2), and complementary sequences for the fiducial mark (FM_comp) on the flow cell surface for an annealing and incubation cycle. This blocked nucleotides that are not relevant for our experiments. We imaged at this step as a background value. We then iterated through the following concentrations of DFAME for subsequent analyses: 0.01, 0.1, 1, 5, 10, 25, 50, 100, 250, and 500 µM. Between each concentration, we washed and incubated the flow cell with 1x Lettuce buffer for 5 min to remove the target and collect new background images. Once the experiment was complete, the flow cell was reset by removing all other reagents by flowing in Illumina PR2 buffer and then stored in 1x IDTE buffer at 4 °C between experiments.

### Functional characterization data processing

As described in Gidi et al. (2025), the images from the instrument were aligned with the FastQ files from the MiSeq. The intensity of each cluster was extracted from the images for each condition and stored with the sequence information in a pandas data-frame that served as the starting point for our analysis (Data link 2). First, the data were filtered to remove control sequences by only keeping sequences that contain the poly(T)10 linker present in our libraries. These data were filtered for sequences that have >95% accuracy according to their Q-score. Next, we normalized each cluster to its relative size by dividing by the intensity of the FAC-Cy3 image. We subtracted the background signal from each cluster by subtracting the preceding cycle’s buffer condition intensity value for each concentration and filtering out any negative values. We aggregated the data for each sequence by taking the mean and standard deviation of the intensity values for each concentration across all replicates of that sequence at each concentration. Finally, we identified each sequence’s mutations and collected them in a string in the data-frame.

For the t-SNE analysis, we fed our sequences and the mean intensity values and standard deviations for each concentration into the algorithm^21^. We plotted the data based on the two-Gaussian mixture model (GMM) outlier analysis and replicate number (Fig. S3). Having identified outliers that were not related to the number of replicates, we vector-quantized clustering into 20 k-means clusters and found that VQ-cluster 19 seemed to capture the outliers well. We then used principal component analysis (PCA) to determine the characteristic features of the data. We looked at the first five features and noticed significant homogeneity in the last four, whereas the first feature (PC0) seemed to capture the outlier behavior (Fig S4). Upon plotting projections of the average centered-log-binding feature for each cluster, we noticed a significant deviation from the mean at the lowest tested concentrations of DFAME for VQ-cluster 19. This led us to believe that this feature captured the binding profile of the mutants. We then created a 2D single and double mutant heatmap of the PC0 projection, which we used to determine ideal locations for a split site based on high levels of deviation from the mean of the PC0 projection.

### Surface-based characterization and validation

The Lettuce aptamer was originally selected and optimized in solution phase. To assess whether it retained the same binding affinity and behavior when modified, we ordered variants of the Lettuce aptamer with either a poly(T)10 linker or a biotinylated poly(T)10 linker. For initial characterization, we mixed each of these aptamers (as well as unmodified Lettuce) with 1 µM DFAME in 1x Lettuce buffer in 150 µL aliquots and tested their response at concentrations of 0.01, 0.1, 1, 10, and 100 µM DNA. We denatured the aptamers in in the presence of DFAME in a thermocycler with the lid temperature set to 105 °C and the block temperature set to 95 °C for 3 min, followed by annealing for 10 min at 21 °C. We then transferred 50 µL of each sample into a well of a half-area 96-well plate in triplicate and measured the fluorescence on the plate reader with gain set to 55, excitation at 540 nm, and emission at 630 nm. We fit the equations to the specific binding Hill slope equation from GraphPad Prism: 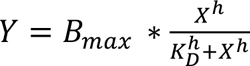.

We next tested the performance of the wild-type Lettuce aptamer with a biotinylated ploy(T)10 linker on streptavidin-coated MyOne Dyna C1 magnetic beads at an aptamer concentration of 1 µM and a bead concentration of 0.25 mg/mL. We incubated the aptamer and beads together rotating for 1 hr at room temperature. In addition to the Lettuce sequences, we incubated the beads with a random DNA control (Random_DNA, Table S1) of similar length to serve as our background measurement. We then washed away unbound aptamers three times with 1x Lettuce buffer and incubated with various concentrations of DFAME for 30 min. After the incubation, we immediately ran the samples through the flow cytometer using the PE-Texas Red channel for excitation and emission at 561 nm and 615/20 nm, respectively. Surface-based measurements are reported as background-subtracted relative fluorescence units, where each point has the background fluorescence value subtracted from that of the random DNA control: F_sample_ – F_NSB_ = F_Background_subtracted_.

After performing our clustering analysis, we characterized the binding of surface-bound versions of a variety of cluster 19 (WT, C20T, T15A, T15A_C20T, T15G, T28A) and non-cluster 19 aptamers (A14T_A21T, C20T_G45A, G26A_G44T, G22C, G19A_A21T, G16C_G24C, A14G_G22T) that had been modified with biotinylated poly(T)10 linker using the same bead-based assay.

### Split Lettuce aptamer design and testing

Split aptamers were ordered from IDT along with reverse-complement DNA sequences for appropriate locations of the RNA target (SARS_split and H5Nx_split sequences in Table S1). We tested them in 1x high-Mg2+ Lettuce buffer (40 mM HEPES, 120 mM KCl, 5 mM MgCl2), unless otherwise noted. All reagents were mixed at concentrations of 1.5 µM of each split aptamer sensor fragment, 2 µM DFAME or DFHBI-1T, and 0.5 µM RNA target before being denatured in a thermocycler by heating to 70 °C with lid temp set to 105 °C for 5 min and then annealed by resting at room temperature for at least 10 min^9^. 50 µL of each sample was transferred from PCR tubes to the wells of a half-area, opaque CoStar 96-well plate and fluorescence measurements were taken on the microplate reader. Measurements with DFAME were taken with excitation at 545 nm and emission at 630 nm. Measurements with DFHBI-1T were taken with excitation at 430 nm and emission at 540 nm. We defined sensor performance as the fluorescence signal-to-background ratio, where background was measured from the sensor in the absence of RNA target and signal was measured from the sensor in the presence of RNA target. To determine the observed, model-derived K_D_ of each split design with DFAME, we mixed each split aptamer sensor fragment at concentrations of 1.5 µM with 2 µM DFAME and 0, 0.1, 1, 10, 100, 500, 750, or 1000 nM RNA target. To determine the observed, model-derived K_D_ of each split design with DFHBI-1T, we mixed each split aptamer sensor fragment at concentrations of 1.5 µM with 2 µM DFHBI-1T and 0, 0.1, 1, 10, 100, 500, 1000, or 2000 nM RNA target. We transferred 50µL of reaction to a half-area, opaque CoStar 96-well plate. We then fitted the data to our derived binding model for the multistate system (Fig. S12). For the full derivation please see the supporting information. From this model, we report the fitted equilibrium constant for chromophore binding to the fully formed split-aptamer/RNA complex as an observed, model-derived K_D_. We use this value as a comparative metric for evaluating sensor behavior across constructs and targets (Fig. 4, S13, and S14). Because some dose-response curves do not fully reach saturation over the tested concentration range, these values should be interpreted as model-derived comparative estimates rather than definitive thermodynamic constants.

### Kinetic Monitoring for Split Sensors

To monitor the kinetic response for each split design, we started by mixing 1.5µM of each split half with 10µL of the concentrated RNA target in 1x high-Mg2+ Lettuce buffer in a total volume of just under 160µL. We then incubated these using a thermocycler with the lid temperature set to 105°C and the block set to 70°C for 5 minutes. We added the appropriate chromophore to a final concentration of 2µM in 160µL. We quickly separated the samples into 50µL for each well in a half-area, opaque CoStar 96-well plate and immediately started measurements on the platereader.

Measurements were taken every 70s for up to 70 minutes. We used the same excitation and emission wavelengths for endpoint fluorescent measurements as mentioned above. We then fit this data using the derived binding model. We used the maximum fluorescent values as the points for the dose response curves and fit those curves with the derived paraments from the binding model. We report the observed, model-derived K_D_ for chromophore binding to the fully formed split-aptamer/RNA complex as a comparative metric for evaluating sensor behavior across constructs and targets (Fig. S13 and S14).

## Statistics

All statistics were performed within the GraphPad Prism software using the built-in 2wayANOVA analysis for comparison between samples, where ****P<0.0001, ***P=0.002, **P=0.0021.

## Data availability

All data from these experiments will be made available at the time of publication. Requests for the data files made be made to the corresponding author.

## Code availability

All code for the analysis of the sequence and functional screening is made available at the following link: https://github.com/Soh-Lab/rationally_designed_lettuce.git.

## RESULTS AND DISCUSSION

### Mutational analysis of the Lettuce aptamer to identify candidate split sites

Inspired by the design and utility of split aptamer sensors, we sought to determine and characterize unconventional split sites for the Lettuce aptamer, using mutational analysis to guide split site location (Fig. 1A). We began our study by sequencing and characterizing a mutant library of all single and double mutants in the target-binding loop region of the Lettuce aptamer, followed by computational analysis of the data, split site identification, and split aptamer characterization (Fig. 1B). We took advantage of our recently developed MAPA platform^19^, which allows for the sequencing and simultaneous functional characterization of many thousands of individual mutant sequences on the flow cell of a benchtop sequencing instrument. MAPA is equipped with excitation lasers at 532 nm and 660 nm, making DFAME, which has an excitation maximum near 545 nm and emission from approximately 610–630 nm, the most compatible chromophore for flow-cell screening.

We were interested in exploring unconventional split site locations, and as such looked to mutate bases in the binding region of the aptamer. Passalacqua and colleagues recently used X-ray crystallography and cryo-electron microscopy (cryo-EM) to characterize the structure of the Lettuce aptamer in complex with DFHBI-1T, DFHO, and DFAME^14^, revealing a four-way-junction and a variety of noncanonical base interactions (Fig. S1). We used this structural information to guide the design of our mutant aptamer library, creating all single and double mutants for bases 7–28 and 43–47, as these nucleotides are involved in the target-binding loop region of the aptamer and not the stem regions (Fig. 1C). Notably, these regions also contain nucleotides that contribute to two G-quadruplexes that were shown to be very important for assembly of the full aptamer and its binding to chromophores. These bases would not typically be considered for split site locations, but we wanted to characterize the tolerance of the loop region to mutations as a proxy for split site locations.

The MAPA platform requires a surface-bound aptamer, while Lettuce was originally selected in solution phase, so we tested if the binding properties of the aptamer were retained when surface bound. To mimic the MAPA platform, we used the same linker sequence, length, and added a surface immobilization moiety, biotin. We confirmed that the Lettuce aptamer retained normal function when a poly(T)_10_ linker and biotin functional group were introduced to its 5′ end via a plate-reader assay (Fig. 1D) Lettuce has a previously reported affinity of 1.9 ± 0.2 µM for DFAME^14^, and we measured similar affinities for unmodified Lettuce (WT), Lettuce with a poly(T)_10_ linker (WT-polyT10), and Lettuce with a biotinylated poly(T)_10_ linker (WT-polyT10-biotin) of 2.5 ± 1.2 µM, 1.7 ± 0.4 µM, and 2.7 ± 1.0 µM, respectively. To further confirm that the biotinylated poly(T)_10_-modified aptamer retained its DFAME affinity when coupled to the surface of streptavidin-coated beads, we incubated both our Lettuce-coated beads and beads coated with a random DNA control with DFAME. We used the control beads to subtract out background fluorescence due to nonspecific binding. We were able to fit our binding data to the Hill slope equation and saw that surface-bound Lettuce had a slightly enhanced K_D_ of 0.098 ± 0.032 µM (Fig. S2), indicating that it should be compatible with surface-based measurements on the MAPA platform.

Using an Illumina MiSeq benchtop sequencer, we generated a flow cell containing hundreds of replicated clusters of individual mutant sequences in order to screen binding behavior of each mutant to DFAME. After its generation, we prepped the flow cell for characterization experiments by blocking DNA that was not part of the mutant library via complements to inhibit any potential interactions (Fig. 1E). We incubated the flow cell containing the mutant library with several concentrations of DFAME in the range of 0.01–500 µM and imaged the fluorescence response at each concentration (Fig. 1F). We utilized DFAME fluorescence intensity as a measurement of binding affinity for each mutant. We then extracted the individual cluster intensities from each image and mapped it back to the x-y coordinates from the sequencing run, enabling us to link fluorescence intensities with the corresponding sequence information and understand the impact of various mutations in the loop region of the aptamer on DFAME binding.

We subsequently evaluated the image data from our MAPA experiment in order to identify promising candidate split sites for further experimental characterization. We extracted the intensity values from each image file as described in our previous work^19^, with the intensity value for every cluster recorded for each condition and mapped to the sequence at the appropriate spatial coordinates. This allowed us to measure the response for each mutant at any given concentration of DFAME as well as its background fluorescence in the absence of DFAME. We identified all 3,004 mutant sequences in our pool and aggregated and normalized the data by averaging the intensity value across all replicate clusters for every sequence at each concentration.

### Computational analysis of the mutational data

We vector-quantized (VQ) each mutant sequence’s binding profile to determine which mutants retained binding. To better visualize the variations in our data from our MAPA platform, we took the logarithm of each sequence’s intensity-vs-concentration profile, after appropriate background-subtraction, and subtracted from each profile the average logarithmic profile from all sequences. We denote these as centered-log-binding features. The mean-subtracted logarithmic intensity-vs-concentration features were dimensionally reduced into two-dimensional space using the t-distributed stochastic neighbor embedding^21^ (t-SNE) algorithm, and vector quantized using k-means clustering into 20 classes (Fig. 2A). We confirmed that the clustering was not artificially based on the number of replicates for each sequence but rather based on the similarity in behavior between the various sequences (Fig. S3A). In contrast, we independently found an outlier population of sequences by fitting all centered-log-binding features into a two-Gaussian mixture model (GMM) (Fig. S3B). The outlier class was found to overlap strongly with the VQ-cluster 19, which contained 127 out of the total 3,004 sequences, comprising just over 4% of the mutant library.

**Figure 2.**
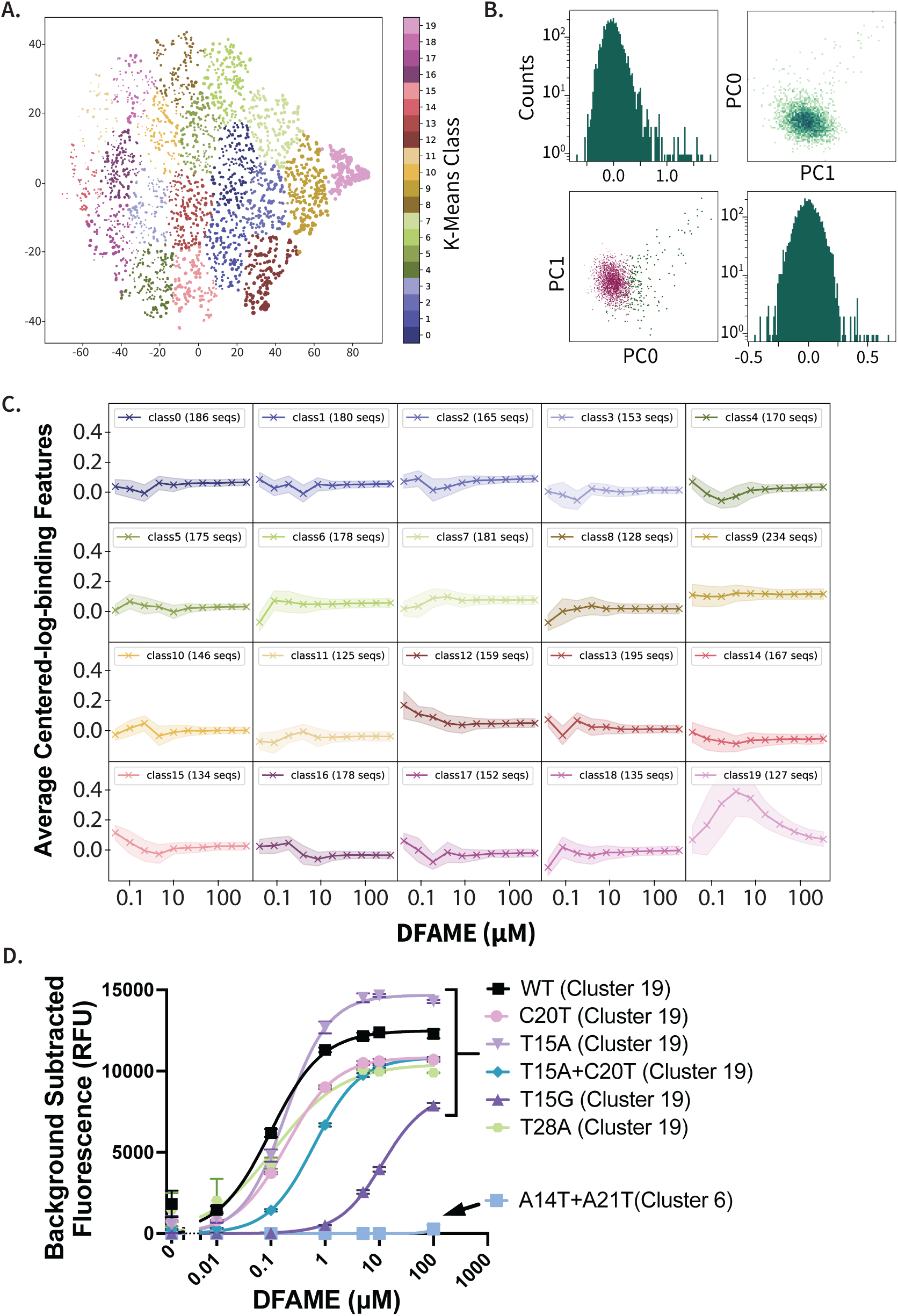
Analysis of the mutant Lettuce library. A) Forced clustering analysis of sequences using t-SNE. Vector-quantized cluster 19 (pale pink) represents the outliers that maintain binding. B) Principal component analysis (PCA) of the entire dataset. PC0 captures the only major deviations in behavior from the average, and we hypothesized that this indicates binding behavior. C) Average centered-log-binding features within each class, with a noticeable increase in deviation from the mean at lower DFAME concentrations indicating binding for VQ-cluster 19. D) Binding curves for WT Lettuce, five cluster 19 sequences, and a non-binding cluster 6 mutant. For each point, we subtracted the background fluorescence value from the random DNA control. N = 3, error bars represent standard deviations, and curves are fit to specific binding Hill slope.

We next studied how VQ-cluster19 varied from the other clusters. Using principal component analysis (PCA) on the MAPA data, we found that PC0 best represented the average centered-log-binding features (Fig 2B; Fig S4). The average centered-log-binding features in VQ-cluster 19 showed a substantial deviation from the mean at lower concentrations of DFAME (Fig. 2C), which was absent in the other clusters. We hypothesized that this deviation was indicative of binding behavior. To confirm this, we randomly selected five cluster 19 sequences (C20T, T15A, T15A + C20T, T15G, and T28A) and eight sequences from other clusters (1, 2, 6, 8, 9, 10, 11, and 18) and studied their binding behavior via a flow cytometry assay in which we coupled biotinylated poly(T)_10_ aptamer sequences onto streptavidin-coated beads and then incubated these (and our random DNA control beads) with various concentrations of DFAME (Fig. S5A). We then made binding curves for WT Lettuce and these various selected sequences with background-subtracted values. For the VQ-cluster 19 sequences, we observed a tight range of binding and characteristic sigmoidal binding curves (Fig. 2D). VQ-cluster 19 sequences responded to DFAME with K_D_s ranging from 0.1–11.1 µM, whereas none of the non-cluster 19 sequences tested showed binding to DFAME (Fig. 2D; Fig. S5B). The distinctive binding behavior of VQ-cluster 19 sequences indicated that we could use the locations mutated within this sequence cluster as split sites, since perturbations at these locations do not seem to interfere with DFAME recognition.

### Identifying candidate split sites in Lettuce that preserve aptamer-ligand binding

Having homed in on a clustering feature predictive of DFAME binding, we next applied this parameter to identify specific nucleotides where a split site could be introduced. To begin, we visualized all single and double mutants in our library with a heatmap reflecting the magnitude by which their PC0 value deviates from the mean (Fig. 3A). Darker blue squares on this heatmap indicate a larger deviation from the mean PC0 (*i.e.,* the binding feature), indicating a K_D_ similar to that of the WT Lettuce aptamer and suggesting that mutations at that site are well-tolerated and could offer useful split sites. Most mutations resulted in much weaker or no binding at all to DFAME, but we did identify a few pockets of mutations that maintain robust binding. We specifically focused on sequences with a single mutation at T28 as well as double mutants containing T28, T15, G16, and C20 (Fig. 3A). Although we observed other areas with tolerated mutations, these nucleotides were of particular interest due to their location within the experimentally defined aptamer structure. T15 and T28 form a non-canonical base interaction just before the start of a second stem-loop region in the aptamer (Fig. S1B). Bases 35–38 from the end of this same stem were removed to generate the original split Lettuce aptamer 5.1.5^9^. G16 is immediately adjacent to T15-T28 and stabilizes this interaction through a noncanonical triplex with G13 and C27 (Fig S1B). C20 was of particular interest because of its location within the binding pocket of the aptamer; this position was previously identified as an essential nucleotide that forms a hydrogen bond with one of the magnesium ions that stabilizes the chromophore-binding site of Lettuce^14^ (Fig. S1B).

**Figure 3.**
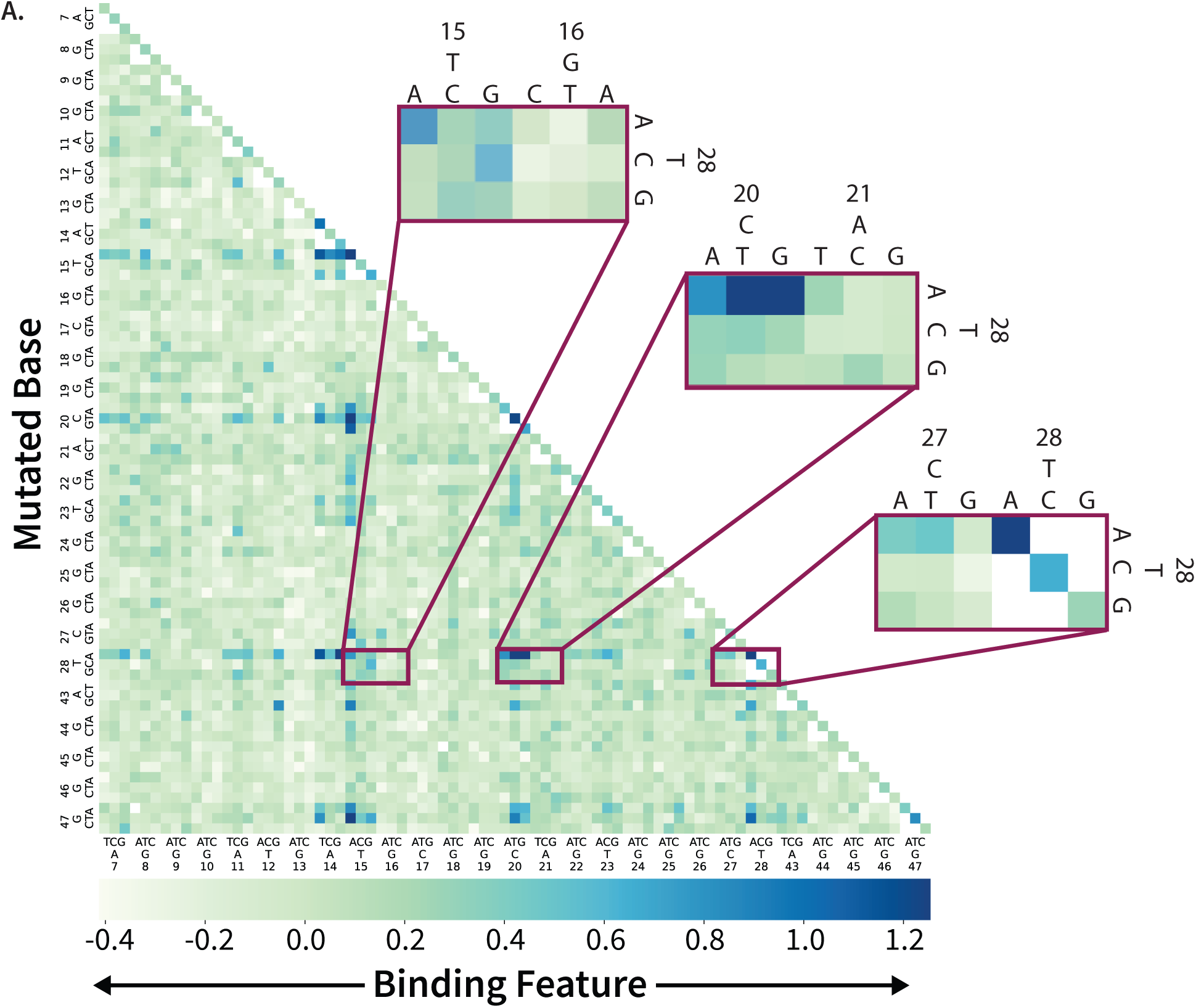
Mutant heat map predicting the positions best suited for splitting the aptamer. A) Single and double mutant heatmap showing the binding feature (PC0) projection for all mutants. Magnifications show sites where mutations are especially well-tolerated (dark blue squares), representing promising split sites. Each split site is represented as a 3D PyMOL rendering in the Supplementary Information, Figures S6–S9.

We developed three split Lettuce aptamers with cleavage sites at each of these three locations: T15/G16 (Fig. S7), C20/A21 (Fig. S8), and T28/T29 (Fig. S9). It is important to note that these locations would not have been identified without mutational analysis because they appear to be in potentially important binding regions within the primary loop of the aptamer, and conventional design principles encourage creating splits far from the known or hypothesized binding pocket^13^. Two of these split sites (T15/G16 and C20/A21) are situated close to G-quadruplexes, which is a counterintuitive choice for a split site, as conventional wisdom suggests that the resulting fragments would not be able to recover chromophore binding when brought back together. Nonetheless, our mutational data suggested that these mutations were well-tolerated, and we proceeded to experimentally assess the binding performance of the split sites predicted in our analytical process.

### Validating the utility of predicted aptamer split sites for sensor development

We experimentally confirmed our split site predictions by generating a split-aptamer Lettuce sensor for an RNA fragment derived from the SARS-CoV-2 viral N gene that substantially outperforms a previously developed sensor based on Lettuce split aptamer 5.1.5^9^ for that same RNA target. We selected this construct as the benchmark because it was previously optimized through extensive mutational and structural studies and used for SARS-CoV-2 RNA detection. For each of the split aptamer sensor designs identified in the previous section (T15/G16, C20/A21, and T28/T29; Fig. 4A), we tagged the 5′ end of the first fragment with a 15-nucleotide (nt) RNA recognition sequence and the 3′ end of the other fragment with a 21-nt RNA recognition sequence, replicating the recognition elements of the previous sensor^9^. We then incubated these fragments with a 36-nt viral RNA fragment to assess the extent to which target binding could bring the two halves back together and produce a fluorescent readout in the presence of DFAME (Fig. 1A). We annealed 1.5 µM of each fragment with 2 µM DFAME and 0.5 µM RNA target for 5 min at 70°C before cooling at 21 °C for 10 min and measuring fluorescence on a plate-reader. All three split aptamer sensors recovered DFAME binding and produced a significant increase in fluorescence in the presence of SARS-CoV-2 RNA, but the T28/T29 split exhibited especially enhanced fluorescence when compared to other designs (Fig. 4B). We observed a 5-fold target-dependent fluorescence enhancement with this construct, compared with the 1.3-fold enhancement observed under the same conditions for the 5.1.5 split-Lettuce sensor, the best split Lettuce aptamer design identified to date. In contrast to the originally published sequence from VarnBuhler *et al.*^14^, which improved split performance with a truncated stem, we utilized the sequence from Passalacqua *et al.*^14^, which included an extra 5-bp stem for crystallization purposes, due to its solved experimental structure. We determined that our T28/T29 split sensor had an observed, model-derived K_D_ of 553.9 nM for SARS-CoV-2 RNA (Fig. 4C).

**Figure 4.**
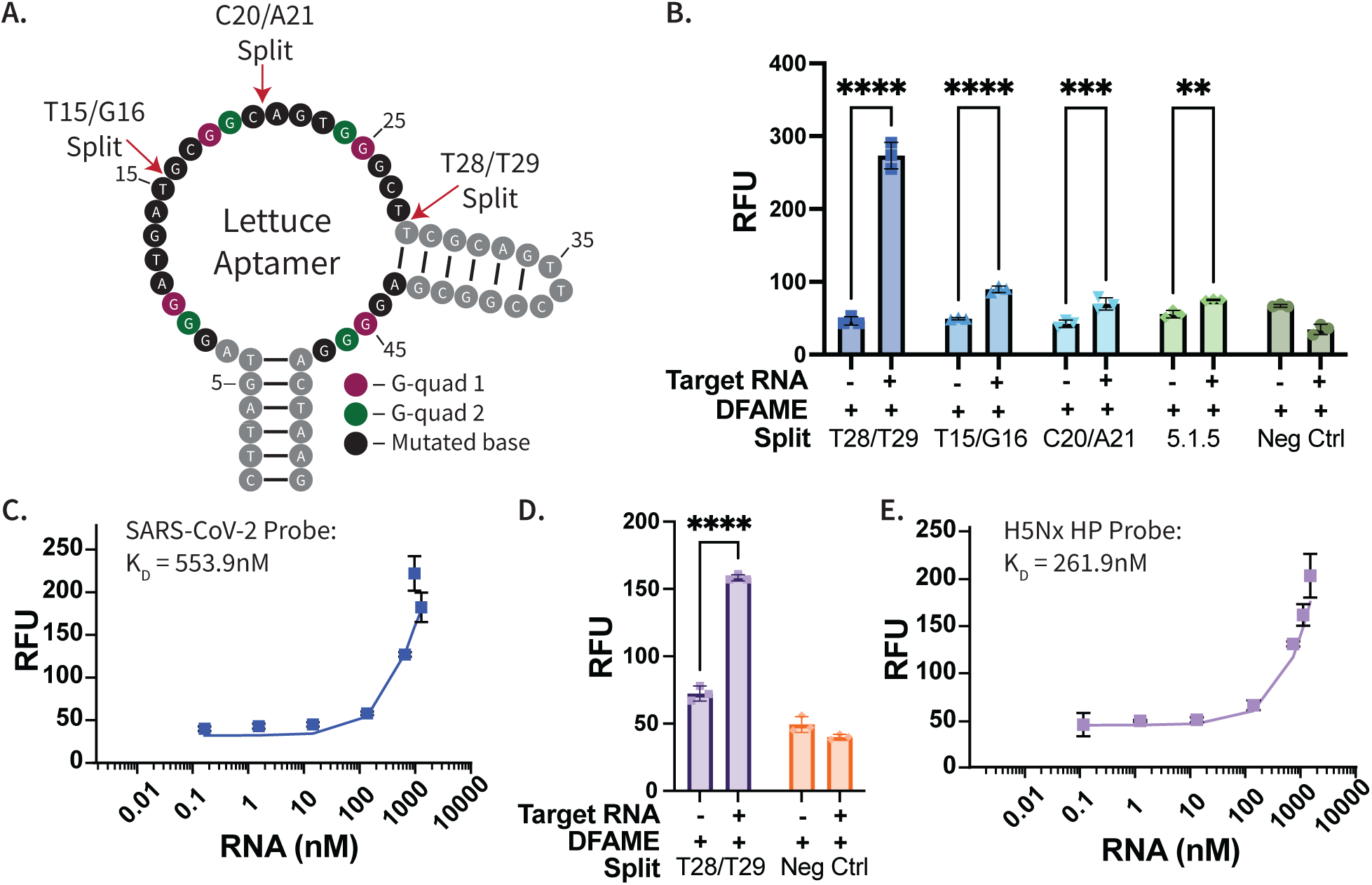
Testing split Lettuce aptamer sensors for SARS-CoV-2 and H5Nx RNA targets. **A**) Diagram of the Lettuce aptamer with the three newly identified split sites indicated, see Fig. S6 for a representative 3D structure. **B**) Fluorescence response of all three split aptamer designs in the presence or absence of SARS-CoV-2 RNA. **C**) Binding curve of the T28/T29 split sensor against SARS-CoV-2 target RNA. **D**) We also designed a T28/T29 sensor for H5Nx high-pathogenicity avian influenza and measured fluorescence in the presence and absence of viral RNA. **E**) Binding curve of the T28/T29 split aptamer sensor against H5Nx HP RNA. N=3, error bars represent standard deviation. ****P<0.0001, ***P=0.002, **P=0.0021. Curves were fit using the multistate binding model described in the Supporting Information.

While testing our split sensor designs, we noticed a large dependence on the magnesium concentration in the buffer, such that the T28/T29 split construct did not produce a signal unless the [Mg^2+^] was five-fold higher than the concentration used in earlier studies with unsplit Lettuce (Fig. S10). Based on these results, we used 5 mM MgCl_2_ in the buffer for all split sensor experiments. Additionally, we evaluated the optimized split constructs with DFHBI-1T under solution-phase conditions. These experiments showed that the improved response of the T28/T29 split sensor was not specific to DFAME under the tested conditions (Fig. S11), which was expected given the similar binding affinity described previously^14^.

The greatly improved performance of the T28/T29 split sensor relative to previously developed split Lettuce aptamer sensors encouraged us to explore its generalizability. We used the same T28/T29 split Lettuce aptamer to design a sensor with a 27-nt RNA recognition sequence that enables the sensor to accurately discriminate high versus low pathogenicity strains of avian influenza H5Nx identified in Eurasian fowl populations^20^. The 5′ end of the first fragment was tagged with a complementary 12-nt target recognition sequence, and the 3′ end of the other fragment was tagged with a complementary 15-nt target recognition sequence. As a target, we chose a short 27-nt RNA fragment of RNA from contemporary clade 2.3.4.4b H5 HP avian influenza. Although the H5Nx target RNA was 9 nt shorter than the SARS-CoV-2 target RNA, the T28/T29 split-Lettuce sensor still produced a target-dependent fluorescence increase in the presence of H5Nx RNA, supporting that our split architecture can be successfully applied to other RNA target (Fig. 4D). Our split construct exhibited an observed, model-derived K_D_ of 261.9 nM for H5Nx (Fig. 4E). However, the H5Nx sensor exhibited a lower signal-to-background ratio and slower kinetic response than the SARS-CoV-2 sensor, indicating that target-sequence and probe-design parameters can still influence sensor performance. The shorter probe length likely accounts for both the decreased signal-to-background ratio of this construct and the slower kinetic response (Fig. S13). These results suggest that the identified split site provides a useful starting architecture, while more target-specific probe optimization may be needed to maximize performance for individual RNA sequences.

## CONCLUSION

In this work, we present a large-scale mutational analysis of the Lettuce aptamer that revealed unconventional split sites within the loop region of the aptamer that yield sensors which outperform existing constructs whose design was informed by conventional wisdom or intuition about appropriate split-site locations. Our MAPA platform enabled the high-throughput analysis of 3,004 single and double mutants of the target-binding loop region of the aptamer. Using t-SNE and PCA, we identified sites within the Lettuce sequence that could tolerate mutations and preserve DFAME-binding capabilities. Based on these tolerated mutations, we identified three especially promising novel split sites along the loop of Lettuce. All three functioned effectively when incorporated into a SARS-CoV-2 RNA sensor, but one split site, T28/T29, showed greatly increased performance relative to other identified sites and yielded a substantially improved Lettuce-based split sensor for this target. Using this same split site, we also developed an effective sensor for high-pathogenicity H5Nx avian influenza RNA.

Critically, the top-performing split site identified in this work would not have stood out as a good choice based on conventional rational design paradigms and would therefore have been overlooked without our mutational analysis. Thus, our work showcases the power of data-driven mutational analysis combined with insights from the 3D structure of the aptamer. Although this study demonstrates the utility of MAPA-guided split-site identification for the fluorogenic DNA aptamer Lettuce, additional aptamer systems will be needed to establish the broader generality of this approach. Importantly, MAPA is not intrinsically limited to fluorogenic aptamers; rather, the platform requires that target binding be coupled to a fluorescence signal. For non-fluorogenic aptamers, this could in principle be achieved using a fluorescently labeled target or a fluorescently labeled competitive strand. However, fluorescent labeling may perturb the native aptamer–target interaction by altering affinity, specificity, folding, or binding kinetics, and each labeling strategy would therefore require experimental validation. Many fluorescent RNA aptamers have been reported, such as Broccoli, Spinach, or Mango. Other groups have used flow cell based technologies to study RNA and peptides, but we have not yet adapted our MAPA platform to be able to accommodate RNA aptamer studies. Furthermore, we were able to take advantage of the experimentally-solved 3D structure for the ligand-bound Lettuce aptamer in this work to guide our mutational analysis and help explain the differences in split site performance. However, only a few such structures have been solved to date for small-molecule-binding DNA aptamers^14,22–28^. Fluorogenic RNA aptamers such as Broccoli^29^, Spinach^30^, and Mango^31^ may also be attractive targets for future MAPA-based split-site analysis. In addition, other groups have demonstrated flow-cell-based approaches for studying RNA and peptides^32–35^. However, we have not yet adapted our MAPA implementation for RNA aptamer studies. To our knowledge, no other fluorogenic DNA aptamers with commercially available chromophores have been reported. As more structures are solved or more fluorogenic DNA aptamers are reported, we believe it will be highly valuable to replicate this approach with a wider range of targets and sequences as a step toward identifying generalizable, evidence-based principles for the rational design of high-performance split-aptamer sensors.

## ASSOCIATED CONTENT

### Supporting Information

The following files are available free of charge.

Detailed DNA and RNA sequences used in studies, PyMOL figures of Lettuce, binding curves of Lettuce and mutants on bead surfaces, expanded PCA and GMM plots, magnesium dependence studies, and DFHBI-1T studies (PDF)

## AUTHOR INFORMATION

A.M.A and H.T.S conceived the project. H.T.S. supervised the project. A.M.A. completed all experiments and data analysis. N.D.L. provided code and analysis for computational methods. Y.G. and L.A.H. developed MAPA, trained A.M.A. and E.B.P and helped with experimental set up. A.M.A., E.B.P. and H.T.S. designed experiments. A.M.A., M.E., and H.T.S wrote the paper. All authors have given approval to the final version of the manuscript.

## ACKNOWLEDGMENT

The authors would like to thank G. Maddocks and L. Zheng for their thoughtful discussions of the work and E. Bagley and N. Torres for lab management support.

## ABBREVIATIONS

MAPA: massively-parallel aptamer performance analyzer
tSNE: t-distributed stochastic neighbor embedding
PCA: principal component analysis
GMM: Gaussian mixture model
VQ: vector-quantized
GFP: green fluorescent protein
FRET: Förster resonance energy transfer
PAGE: polyacrylamide gel electrophoresis.

